# MECAT: an ultra-fast mapping, error correction and *de novo* assembly tool for single-molecule sequencing reads

**DOI:** 10.1101/089250

**Authors:** Chuan-Le Xiao, Ying Chen, Shang-qian Xie, Kai-Ning Chen, Yan Wang, Feng Luo, Zhi Xie

**Affiliations:** State Key Laboratory of Ophthalmology, Zhongshan Ophthalmic Center, Sun Yat-sen University, Guangzhou 510060, China; School of Data and Computer Science, and Guangdong Provincial Key Lab of Computational Science, Sun Yat-sen University, Guangzhou 510275, China; Southern Regional Collaborative Innovation Center for Grain and Oil Crops in China, Hunan Agricultural University, Hunan 410128, China; College of Plant Protection, Hunan Agricultural University, Changsha, China; School of Computing, Clemson University, Clemson, SC 29634-0974

## Abstract

The high computational cost of current assembly methods for the long, noisy single molecular sequencing (SMS) reads has prevented them from assembling large genomes. We introduce an ultra-fast alignment method based on a novel global alignment score. For large human SMS data, our method is 7X faster than MHAP for pairwise alignment and 15X faster than BLASR for reference mapping. We develop a Mapping, Error Correction and de novo Assembly Tool (MECAT) by integrating our new alignment and error correction methods, with the Celera Assembler. MECAT is capable of producing high quality *de novo* assembly of large genome from SMS reads with low computational cost. MECAT produces reference-quality assemblies of *Saccharomyces cerevisiae*, *Arabidopsis thaliana*, *Drosophila melanogaster* and reconstructs the human CHM1 genome with 15% longer NG50 in only 7600 CPU core hours using 54X SMS reads and a Chinese Han genome in 19200 CPU core hours using 102X SMS reads.

## Introduction

Determining the genome sequence of a species or an individual in a population is one of the most important tasks in genomics^1–6^. *De novo* assembly is a process that reconstructs the genome from sequencing reads without a reference genome^7–10^. While technical advances in next generation sequencing (NGS) have enabled to assemble a genome in significantly lower cost and higher throughput comparing to the first-generation Sanger sequencing^11^, two inherited drawbacks make assembly of a genome from NGS short reads difficult^12–14^. First, NGS reads are only few hundreds base pair in length, which are shorter than the lengths of most repetitive sequences in either microbial or eukaryotic genomes^15–17^. Second, PCR amplification in library preparation causes sequencing biases, resulting in some sequence contexts, such as GC-rich regions, not being sequenced^12,17^. Both drawbacks lead to incomplete, fragmented assemblies^18^. The recently emerged third generation single molecular sequencing (SMS) technologies^19^, such as PacBio single molecule real time (SMRT)^20,21^ and Oxford Nanopore^21–26^, posses two distinguishing characteristics, namely, the long read length and the unbiased sequencing^17,27–29^, which can overcome the deficiencies of NGS^17,27–29^. These two properties together may help better resolve repeats and biased region, and thus obtain high-quality *de novo* genome assemblies^21,30–33^.

The SMRT and Nanopore reads usually have high error rates^34–37^. For example, the error rate of PacBio SMRT reads is generally 13-18%^35,38^. However the errors of SMRT are random and dominated by point insertions and deletions^34^ with no preference on particular genome regions^29^. Both theoretical and practical studies have shown that the SMRT reads can be corrected with high accuracy provided the sequencing coverage is high enough^21^. Therefore, a “correction then assembly” approach has been used by assemble pipelines, such as PBcR^35^, FALCON^39^ and HGAP^21^, for single molecular sequencing reads. In those pipelines, raw noisy reads are first corrected and then fed into an overlap graph based assembler, such as the Celera Assembler^3,35^. Previous practices have demonstrated that the “correction then assembly” approach can reconstruct highly continuous and accurate genome assemblies^35,39^.

Although SMRT sequencing have already been widely used to reconstruct small bacteria and archaea genomes, assembling middle or large size genomes from SMRT reads have suffered from high computational cost in the correction step of “correction then assembly” pipelines^3,38–40^. In PBcR-MHAP pipeline, about 84% computational time is used in read correction step. Recently, with new algorithm advancing, the assembly of *D. melanogaster* from SMRT reads takes only 1060 CPU core hours using PBcR-MHAP, which is dramatically reduced from 631, 000 CPU hours using original PBcR^3,41^. However, it still take a very long time for pipelines, such as PBcR-MHAP and FALCON, to assemble large genomes^42^. For example, it costs 260, 000 CPU core hours for PBcR-MHAP and FALCON to complete a human genome from 54X raw SMRT sequences^3,42,43^.

The high computational cost of SMRT assemble pipelines is mainly due to the all-pair alignment step to determine overlaps between read pairs for the correction. There are two sub-steps in the all-pair alignment step. First, the k-mer mapping based approach is used to identify candidate matched read pair. Then, local alignment is used to determine final matched read pair. Due to highly repetitive nature of biology genomes^44,45^, reads sampled from repetitive regions can lead to a high number of k-mer matches^42–44^, which lead to a lot of excessive^44^ candidate pairs. Meanwhile, the local alignment of two long noisy reads is also slow, even with linear local alignment program, like diff^46^. The local alignment of excessive candidate pairs is the major computational time waste in read correction.

To reduce the excessive matched reads after k-mer mapping, and then reduce the total computational time, we present a novel alignment filtering algorithm based on fast global k-mer scoring. Our filtering algorithm is inspired by the observation that the frequency of repeat subsequences decreases dramatically with their size. Thus, if we can find long matched read pairs, then these alignments can be non-repetitive matches with high confidence. We develop a novel k-mer seed score that is correlated with the overlap size between two reads, and then is able to represent the global matching information. For each read, we can select top matched reads according to their k-mer seed score for the further read correction. Noted, selecting top matched reads based on number of matched k-mers may lead to many non-informative matches since the repetitive region has more matched k-mers. Our global k-mer scoring algorithm allows us to dramatically reduce the number of non-informative matched read pairs, as well as selecting smaller number of informative matched reads for the read correction; both can lead to the significantly reducing of computational cost.

We have also presented a new SMS read error correction method by combining counting-based method and local partial order graph, which can achieve high correction accuracy and high correction speed simultaneously. With those new algorithms, we develop an ultra-fast Mapping, Error Correction and de novo Assembly Tool (MECAT) for SMRT reads. MECAT achieves superior computing efficiency to current assembly pipelines. In particular, MECAT takes only about 7600 CPU core hours to assemble a high quality human CHM1 genome using 54x SMRT data^47^ (CHM1) on a single 32-threads computing node with 2.0 GHz CPU, which is 34 times faster than the current PBcR-MHAP pipeline^3^. The MECAT makes it possible to *de novo* assemble large genome using SMRT reads with the similar computational cost as that the assembling of NGS reads needs.

## Results

### Alignment filtering in MECAT

The initial step of our alignment is also finding the candidate alignments by mapping the k-mers of two blocks^3,40,48^ with size of 1,000 to 2,000 bp. Two blocks are considered matched if the number of matched k-mer beyond a predefined threshold. We find a candidate alignment between two SMS reads or between a SMS read and reference genome if there is at least one block pair between them matched. The k-mer matching based method can filter out random pairs and quickly find seed alignments with high sensitivity^40^. However, a read often aligns to many other reads or many locations in the genome due to highly repetitive nature of genomes^42,44^. Local alignments are needed to find the good matched reads or best matched genome locations^48^. However, the computational cost for local alignments between two long SMS reads or between a SMS read and reference genome is high^40,46^. Meanwhile, most of SMS applications, such as SMS read correction and reference genome mapping, only need limited number of matches^48–50^. To quickly select high quality candidate alignments for the further local alignments, we develop a new pseudo linear global scoring algorithm to filter candidate alignments (Figure 1). Our algorithm works by scoring matched k-mer pair in two steps using distance difference factors (DDF). First, we mutually score the k-mer pair in a selected matched block pair. The k-mer pair with max score is selected as the seed, and then the seed k-mer pair is scored by the matched k-mer pairs in other matched block pairs. The score of the seed k-mer pair is supported by all informative matched k-mer pairs and their interval distance when there is a good alignment between them. Thus, our scoring algorithm integrates the global matching information between two SMS reads or between a SMS read and reference genome. Figure 2a shows that the scores of seed k-mer pairs between SMS pairs grow linearly with their overlapping lengths in four SMS data sets. Therefore, by selecting SMS read pairs with high scores, we can filter out the non-informative candidate alignments. After filtering by global scoring, we have reduced 50% to 70% candidate alignments for further local alignment (Figure 2b). And this makes the alignment tool 2-3 times faster than those without global scoring filtering. The candidate alignments are then further filtered by local alignment using diff program^40,46^.

**Figure 1.**
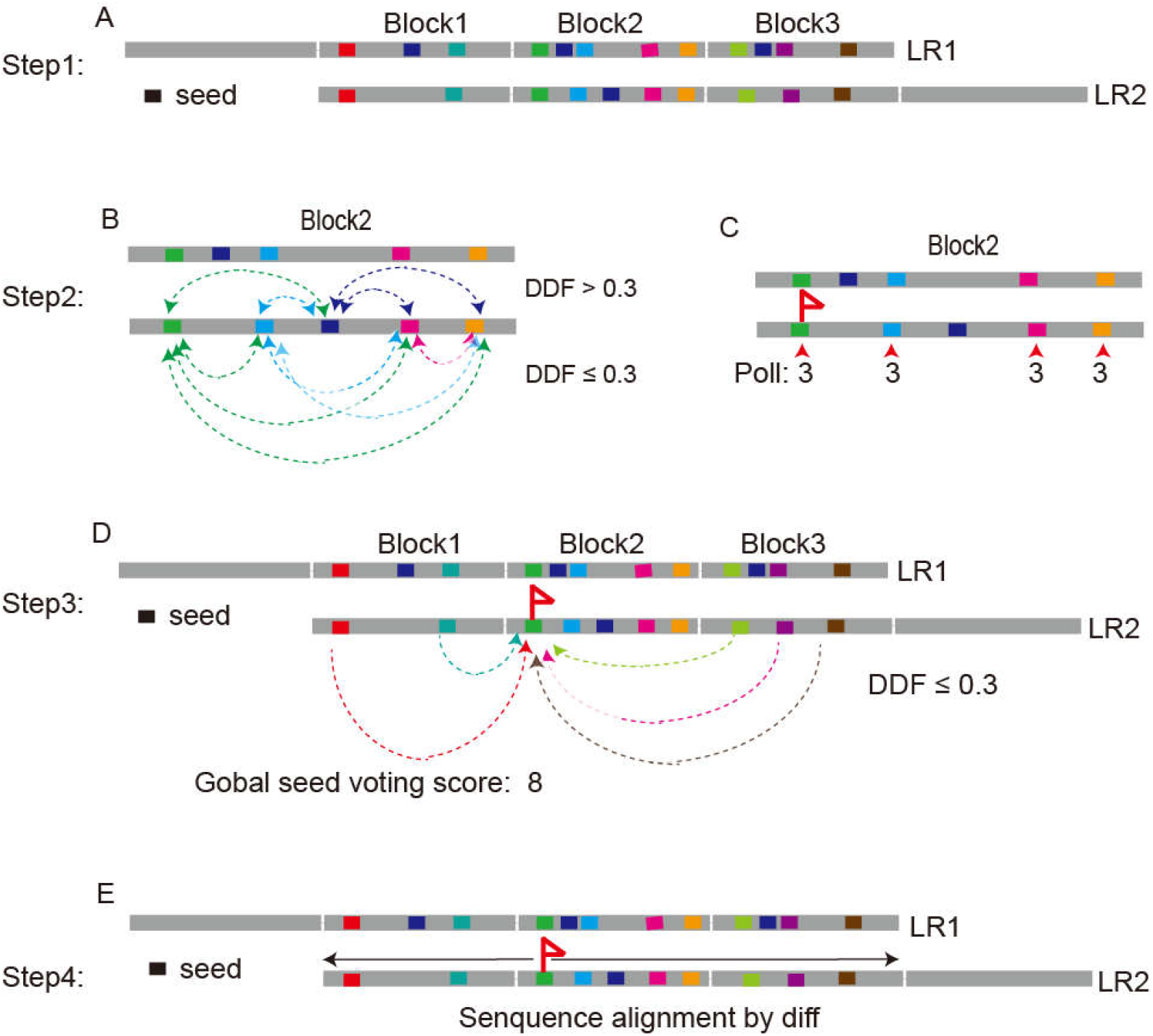
Principle of global scoring algorithm in MECAT alignment. (A) Alignment of k-mers between blocks of two SMS reads. (B) Pairwise scoring using DDF between k-mer pairs in each block pair (Block2 in A as an example). (C) Selecting the seed k-mer pair with the highest score. Random selecting one if multiple k-mer pairs have the same scores. (D) Scoring the seed k-mer pair using k-mer pairs in other block pairs. (E) Aligning two reads from the seed k-mer pair.

**Figure 2.**
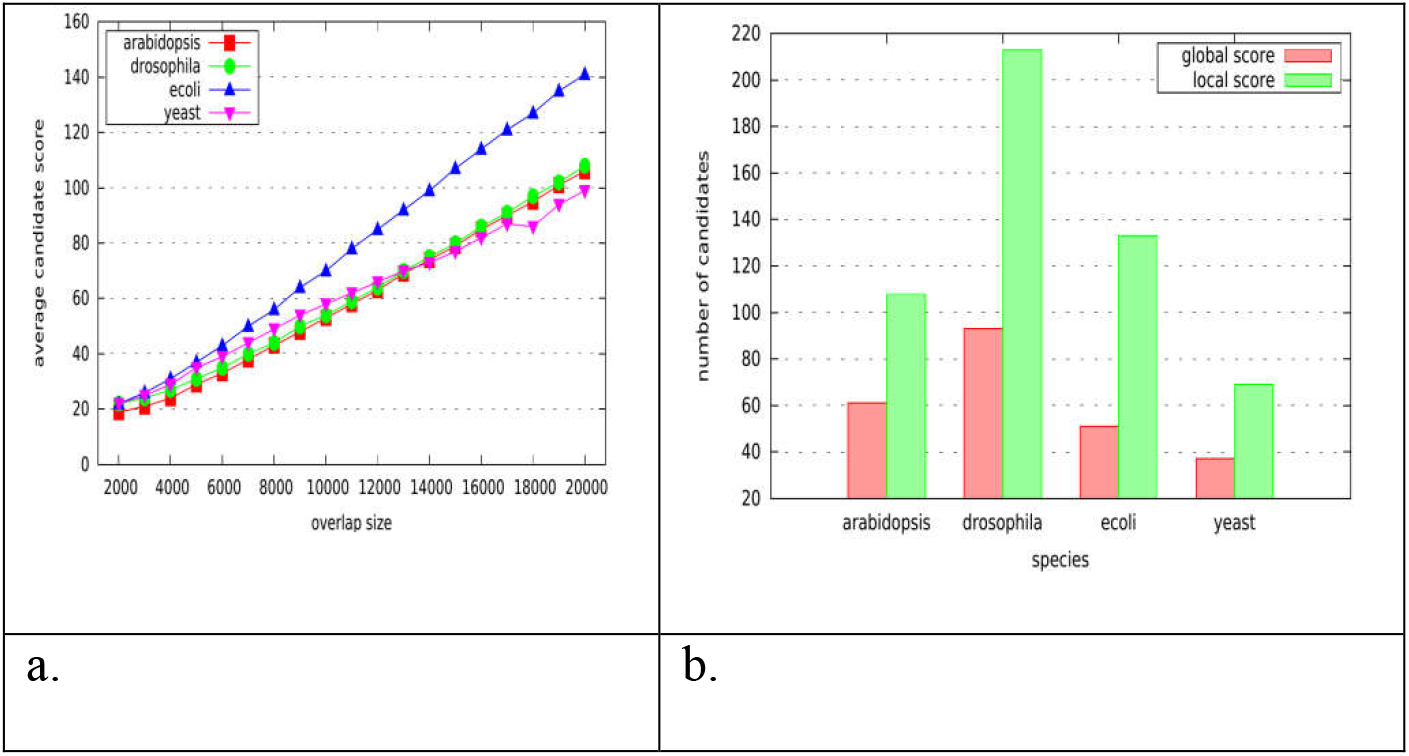
The global scores between SMS read of four model organisms. (a) The relationship between the overlap length of two reads and their global scores. We first extract long reads (length >= 5000bp) from four SMS data sets (E coli, Yeast, A. Thaliana and D. Melanogaster)^47^. We perform pairwise alignment of reads in each data set using MECAT and record the overlap size and its corresponding global voting score of each alignment. (b) Comparison of the numbers of alignment candidates before and after global scoring filtering.

### Pairwise alignment performance of MECAT

To evaluate the performance of MECAT in pairwise alignment of SMS reads, we first compare memory usage and computational time cost of MECAT to those of three widely used SMS read alignment tools, MHAP^3^, BLASR^48^ and DALIGN^40^. We evaluate the alignment tools using five real datasets^47^. As shown in Table 1, the MECAT is faster than all other aligners, except it is slightly slower than DALIGNER^40^ on the E. coli dataset, which is due to computation cost of pre-processing procedure in MECAT. For large human genome, MECAT is 7 times faster than the second best aligner, MHAP-fast, and 15 times faster than DALIGNER. Meanwhile, MECAT uses similar amount of memory that DALIGNER uses, which is only about 1/10 of the amount of memory used by MHAP. Thus, MECAT has used only small amount of memory to achieve fast pairwise alignment.

**Table 1.**
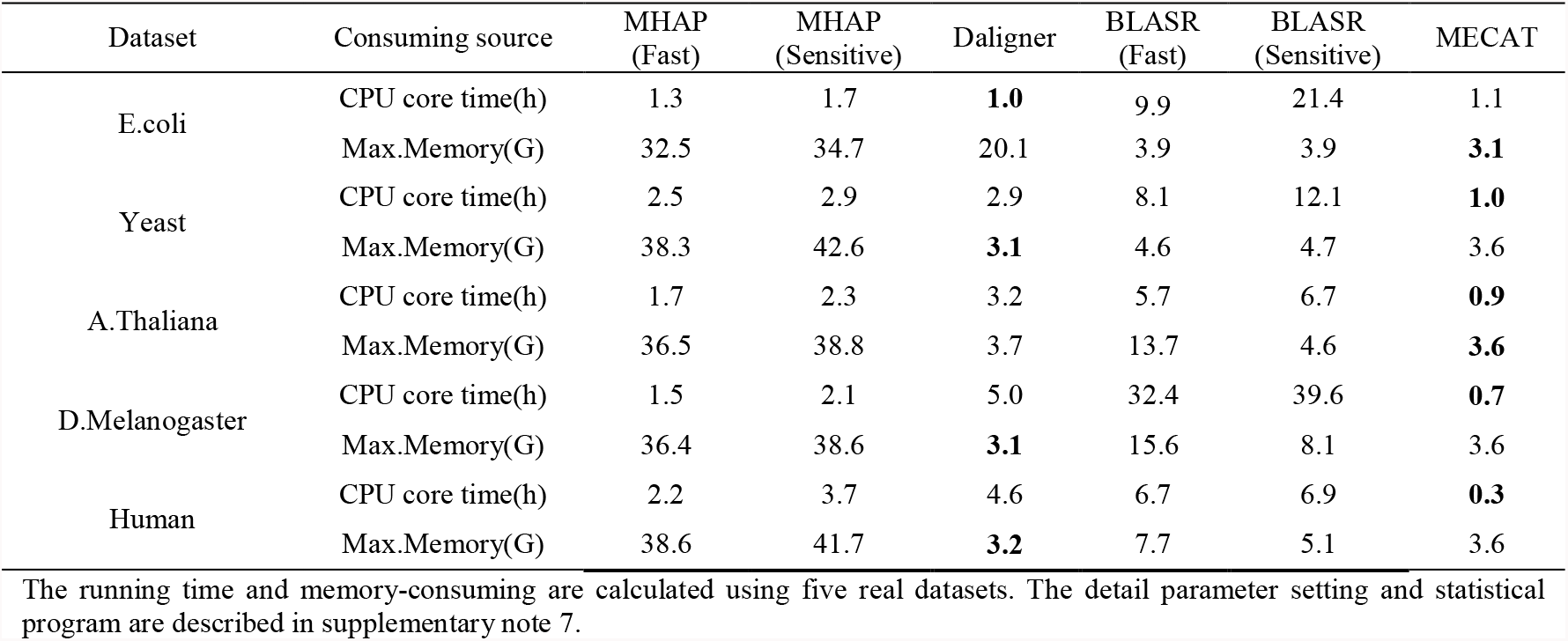
Computing performance of pairwise alignment of SMS reads.

Next, we evaluate the sensitivity and precision of aligners on three simulated datasets, including a 20x coverage E. coli, a 20x coverage yeast and a 5x coverage human chr1 datasets (Table 2). Since we knew the beginning and ending position of each read in reference genome in the simulated datasets, we can calculate the true pairwise overlap relationships between all reads. The sensitivity of an aligner indicates its ability to identify true overlaps and the precision of an aligner indicates the correctness of identified overlaps. The sensitivities of DALIGNER^40^ are the best among four aligners, but its precisions are the lowest. On the other hand, the BLASR^48^ and MHAP^3^ have high precisions, but low sensitivities. Meanwhile, MECAT maintains high precision as well as high sensitivity at the same time. The sensitivities of MECAT are consistently higher than those of both BLASR and MHAP while maintaining similar precisions. Comparing to DALIGNER, MECAT has higher precisions, but lower sensitivities. The precision and sensitivity of DALIGNER become extremely unbalanced for human chr1 data, with only 9.1% precision. These results show that MECAT has achieved a good balance between sensitivity and precision for both small and large genomes.

**Table 2.** Pairwise alignment sensitivity and precision comparison of different aligners.

| Dataset | properties | MHAP<br>(Fast) | MHAP<br>(Sensitive) | Daligner | BLASR<br>(Fast) | BLASR<br>(Sensitive) | MECAT |
| --- | --- | --- | --- | --- | --- | --- | --- |
| E.coli | Sensitivity (%) | 29.7 | 54.4 | **93.7** | 44.5 | 45.1 | 80.3 |
|  | Precision (%) | **99.3** | 98.1 | 90.0 | 98.1 | 89.9 | 96.2 |
| Yeast | Sensitivity (%) | 34.5 | 60.8 | **93.9** | 45.4 | 46.2 | 80.2 |
|  | Precision (%) | 92.2 | 90.2 | 53.9 | **94.9** | 71.7 | 87.6 |
| Human<br>Chr1 | Sensitivity (%) | 37.0 | 63.8 | **93.8** | 41.8 | 46.0 | 66.2 |
|  | Precision (%) | 88.9 | 82.6 | 9.1 | **94.6** | 77.1 | 91.6 |

We define that two read has the correct pairwise relationship if they overlap in reference genome for more than 2k pb. The alignments of each tool result in following three cases: (1) correctly identify the pairwise relationship (C); (2) identify wrong pairwise relationship (I); (3) fail to find the pairwise relationship (U). Then, the sensitivity of the tool is defined as $C/(C+U+I)$ , and the precision of the tool is defined as $C/(C+I)$ .

### Reference genome alignment performance of MECAT

The MECAT can also be used to align SMS reads to the reference genome. We evaluate MECAT with two popular SMS reads reference genome aligners, BLASR^48^ and BWA-mem^49^. We have first compared the aligning time cost using five real datasets (Table 3). For four small genomes (E.coli, Yeast, Arabidopsis, fly)^47^, MECAT is 40 to 85 times faster than BLASR and 20 to 83 faster than BWA-mem. For human genome, MECAT is 15 times faster than BLASR and 5 times faster than BWA-mem. Then, we have compared the sensitivities, precisions and coverages of aligners using 20X simulated SMS data of E. coli, yeast and human genomes (Table 4)^51^, in which we know the read positions on the genomes. Comparing to BLASR^48^ and BWA-mem^49^, MECAT maps slightly less amount of reads to the reference genome, but it map more reads correctly for all three data sets. As a result, MECAT has higher sensitivities, precisions as well as coverages for all three data sets. The results show that MECAT can align SMS reads to the reference genome ultra-fast and maintain high sensitivity and precision. In five real datasets, the mapping overlap rates of three algorithms are as high as 95-99% of the same alignment positions (Supplementary Note 6 and Supplementary Figure 1-5), showing high confidence of MECAT alignment.

**Table 3.** Reference genome alignment speed comparison among MECAT, BLASR and BWA-mem.

| Dataset | Data size | Time (min)/BLASR | Time (min)/BWA-mem | Time (min)/MECAT |
| --- | --- | --- | --- | --- |
| E.coli | 649.4M | 162.2 | 93.7 | 1.9 |
| Yeast | 5.6G | 756.7 | 408.65 | 18.1 |
| A.Thaliana | 36.1G | 2511.7 | 2323.4 | 30.8 |
| D.Melanogaster | 29.3G | 2406.3 | 3244.5 | 38.8 |
| Human | 389.2G | 26940.0 | 9417.5 | 1747.2 |

The running time and memory-consuming are calculated using five real datasets. The detail parameter setting and statistical program are described in supplementary note 6.

**Table 4.** Reference genome alignment sensitivity and precision comparison of different aligners.

| Dataset | Method | SMS reads count | Mapped count | Correct count | Correct mapped length | Precision | Sensitivity | Coverage |
| --- | --- | --- | --- | --- | --- | --- | --- | --- |
| Ecoli | BLASR | 6,634 | 6,634 | 6,606 | 88,207,198 | 99.58% | 99.58% | 99.53% |
|  | BWA-mem | 6,634 | 6,634 | 6,511 | 86,980,847 | 98.15% | 98.15% | 98.15% |
|  | MECAT | 6,634 | 6,634 | 6,633 | 88,592,740 | **99.98%** | **99.98%** | **99.97%** |
| Yeast | BLASR | 17,386 | 17,386 | 17,315 | 231,112,187 | 99.59% | 99.59% | 99.55% |
|  | BWA-mem | 17,386 | 17,386 | 16,921 | 225,918,564 | 97.33% | 97.33% | 97.32% |
|  | MECAT | 17,386 | 17,384 | 17,367 | 231,880,652 | **99.90%** | **99.89%** | **99.89%** |
| Human | BLASR | 4,422,350 | 4,079,186 | 4,040,515 | 53,969,578,429 | 99.05% | 91.37% | 91.31% |
|  | BWA-mem | 4,422,350 | 4,079,186 | 3,925,313 | 52,454,109,629 | 96.23% | 88.76% | 88.74% |
|  | MECAT | 4,422,350 | 4,079,021 | 4,046,195 | 54,073,550,527 | **99.20%** | **91.49%** | **91.48%** |

The aligned positions have one of following three situations, C, I or U, where C indicates alignment to its correct position when the absolute distance of the aligned position and the reference position from pbsim is less than 100bps; I indicates alignment to a wrong position and U indicates fail to be aligned to reference genome. Therefore, sensitivity is defined as $C/(C+I+U)$ , and precision is defined as $C/(C+I)$ . Coverage is defined as the mapped base length / total base length.

### Error Correction Performance of MECAT

Due to high error rates of SMS reads, error correction is an indispensable step when they are used for genome assembly^34–37^. Currently, FC_Consensus, DAGCon^35^ and FalconSense^3^ are three most widely used SMS read error correction methods. DAGCon^35^ represents a multiple read alignment for each read to be corrected as a partial order graph (POG) and find the correct consensus sequence using dynamic programming. DAGCon is accurate but very slow. On the other hand, FalconSense simply corrects the template sequence by counting consistent base alignments. FalconSense is fast but less accuracy. The accuracy of SMS reads is important for genome assembly^35,38^. Here, we develop a new SMS read error correction method by combining the principles from both DAGCon and FalconSense. For regions with consistent matches/deletions without insertion (trivial regions), we use counting-based method. And for other complicate regions, we construct a local POG and solve it with dynamic programming. As the complicate regions are generally less than 10 bases, the local POGs are very small and can be solved very fast.

The first step of read correction is to perform the alignments between the template and the relative reads, which need random access the storage that stores the reads. Both DAGCon and FalconSense store the read in hard driver, which does not support random access. The slow read loading process in DAGCon and FalconSense lead to only 20% CPU usage. To accelerate the correcting process, we load all reads into memory, which supports random access. We encode each base using 2 bits. Thus, the memory occupation of MECAT is about 1/4 of the total read size. Loading reads to memory makes the CPU usage of MECAT over 96%.

We have compared MECAT error correction to FalconSense in PBcR-MHAP^3^ and Canu, as well as FC_Consenses in FALCON using four real datasets. Table 5 shows that the correction running speed from MECAT error correction is 5∼20 times faster than FalconSense and 3∼10 times faster than FC_Consense for four datasets. To evaluating the accuracy of corrected long reads, all corrected long reads were mapping into reference genome by dnadiff program^52^ (Table 5 and Supplementary Note 8). The mapping results show that the accuracies of reads corrected by MECAT are always higher than 99% and are the best in three of four datasets. Specially, for D. melanogaster dataset, the accuracies of other tree method are less than 99%, while the accuracy of MECAT is as high as 99.26% (Table 6 and Supplementary Note 8).

**Table 5.** Comparison of speeds of SMS read correction methods.

| Dataset | FC_Consense from<br>FALCON | FalconSense from<br>PBcR-MHAP | FalconSense from<br>Canu(v1.3) | MECAT |
| --- | --- | --- | --- | --- |
|  | Time(h)/Size/Speed(g/h) | Time(h)/Size/Speed(h/g) | Time(h)/Size/Speed(h/g) | Time(h)/Size/Speed(h/g) |
| E.coli | 0.83/246m/0.29 | 0.36/197m/0.53 | 0.34/158m/0.45 | 0.13/282m/2.13 |
| Yeast | 1.06/603m/0.56 | 1.90/236m/0.12 | 3.22/388m/0.12 | 0.58/716m/1.21 |
| A.thaliana | 59.21/12.1g/0.20 | 41.37/3.10g/0.11 | 30.67/4.09g/0.13 | 4.23/10.05g/2.38 |
| D.melanogaster | 45.21/10.26g/0.23 | 28.57/3.40g/0.12 | 28.37/4.01g/0.14 | 5.75/10.29g/1.79 |

Three algorithms from different pipelines have been tested on the same computer with 2.0 GHz CPU and 512 GB RAM memory by running whole pipeline. Here, Time is the running time of correction process and Size is the corrected sequence size of output result. Speed is the correction speed of correction algorithm and it is defined as Size /Time.

**Table 6.** Comparison of accuracy of corrected reads from of SMS read correction methods.

|  | Raw data | FC_Consenses | FalconSense | FalconSense | MECAT |
| --- | --- | --- | --- | --- | --- |
|  | accuracy | from FALCON | from PBcR-MHAP | from Canu(v1.3) |  |
| E.coli | 90.98 | 99.78 | 99.23 | 99.72 | **99.80** |
| Yeast | 92.70 | 99.55 | 99.01 | 99.48 | **99.57** |
| A.thaliana | 88.24 | **99.57** | 98.60 | 98.79 | 99.36 |
| D.melanogaster | 87.73 | 98.39 | 97.83 | 98.21 | **99.26** |

100M of raw and corrected reads of each datasets are randomly selected to evaluate raw read quality and corrected read quality. Raw reads and corrected reads were mapped on the reference genome by dnadiff software. The mean sequencing error of 100M data was estimated by the in-house R script..

### Assembly performance of MECAT

Generally, there are three steps in the assembly of genome using SMS reads: overlapping SMS reads to selected template reads; correcting the selected reads and constructing the contig using corrected reads^3,35,38^. To test the efficiency and effectiveness of MECAT aligner and error correction in genome assembly, we develop two genome assembly pipelines. In first pipeline, the SMS reads are overlapped and corrected by MECAT, and then fed into Celera Assembler (CA). We call the first pipeline MECAT-CA. In the second pipeline, which is called the MECAT, the SMS reads are also first overlapped and corrected by MECAT. Then, the corrected reads are overlapped by MECAT aligner again and the overlap graph is fed into the “Unitig Construction” module of Canu^42^ (v1.0) to construct the contigs. In both pipelines, we do not perform local alignment using diff program during SMS reads overlapping. We only select the top mapped reads using global scores and feed the mapping information to error correction step. We compare two pipelines to other three SMS assembly pipelines, PBcR-MHAP^3^, FALCON and Canu. We evaluate assembly pipelines using previously released whole genome SMRT reads of five genomes: *E. coli* K12, *S. cerevisiae* W303, *D. melanogaster* ISO1, *Arabidopsis thaliana* Ler-0 and the complete hydatidiform mole CHM1^47^. All the genome assemblies are polished by Quiver^53^ to correct sequencing errors.

Table 7 lists the running time of five pipelines on the same computer. We evaluate total assembly time as well as the running time for read overlap, error correction and contig construction separately. For small *E. coli* K12 genome, the MECAT-CA and MECAT are 3.7 to 5.0, 3.9 to 5.4 and 2.2 to 2.9 times faster than FALCON, PBcR-MHAP and Canu, respectively. For another small *S. cerevisiae* W303 genome, the MECAT-CA and MECAT are 1.5 to 2.1, 3.2 to 4.3 and 3.9 to 5.3 times faster than FALCON, PBcR-MHAP and Canu, respectively. For medium *D. melanogaster* ISO1, the MECAT-CA and MECAT are 7.32 to 14.7, 5.3 to 10.5 and 3.6 to 7.2 times faster than FALCON, PBcR-MHAP and Canu, respectively. For another medium *Arabidopsis thaliana* Ler-0 genome, the MECAT-CA and MECAT are 15.5 to 20.9, 13.1 to 17.6 and 8.2 to 11.1 times faster than FALCON, PBcR-MHAP and Canu, respectively. For large human genome, we are not able to run other assembly pipelines on our single computer, thus we compare our running time to the results of previous paper^3^. Our MECAT-CA is 5.2 and 12 times faster than PBcR-MHAP-fast and PBcR-MHAP-sensitive, and MECAT shows remarkable 24.9 times speedup than PBcR-MHAP-fast and 56.4 times speedup than PBcR-MHAP-sensitive. The larger the size of genome is, the greater the speedups of MECAT are.

As shown in Table 7, the overlapping and correcting steps are the most time consuming steps among the three assembly steps of PBcR-MHAP, FALCON and Canu. And the speedups of MECAT-CA and MECAT are mostly coming from the efficient of MECAT aligner and error correction in these two steps. For small or medium size genomes, the running times of three steps in MECAT-CA pipeline are similar. However, the running time of contig construction became the bottleneck comparing to other two steps for large genome. For human genome, the running time of contig construction step are 3.5 and 39 times longer than those of overlapping and correcting steps in MECAT-CA pipeline. Meanwhile, for human genome, the running time of contig construction of MECAT is only 7.5% of that of MECAT-CA, which make contig construction step not a bottleneck in MECAT, in which MECAT pairwise alignment replaces OverlapIncore program in CA.

**Table 7.** The computational time comparison of different assembly pipelines.

| Genome | Pipeline | threads | overlap time(h) | correct time(h) | correction size | Contig time(h) | Total time(h) |
| --- | --- | --- | --- | --- | --- | --- | --- |
| E.coli | FALCON | 16 | 0.22 | 0.83 | 246M | 0.15 | 1.21 |
|  | PBcR-MHAP | 16 | 0.74 | 0.36 | 197M | 0.15 | 1.29 |
|  | Canu | 16 | 0.20 | 0.34 | 158M | 0.15 | 0.71 |
|  | MECAT-CA | 16 | 0.05 | 0.13 | 282M | 0.15 | 0.33 |
|  | MECAT | 16 | 0.05 | 0.13 | 282M | 0.05 | 0.24 |
| Yeast | FALCON | 16 | 0.72 | 1.06 | 603M | 0.28 | 2.06 |
|  | PBcR-MHAP | 16 | 1.47 | 1.9 | 236M | 0.45 | 4.2 |
|  | Canu | 16 | 1.15 | 3.22 | 388M | 0.67 | 5.11 |
|  | MECAT-CA | 16 | 0.12 | 0.58 | 716M | 0.63 | 1.33 |
|  | MECAT | 16 | 0.12 | 0.58 | 716M | 0.25 | 0.97 |
| A.thaliana | FALCON | 16 | 132.52 | 59.21 | 12.1G | 32.11 | 223.84 |
|  | PBcR-MHAP | 16 | 132.76 | 41.37 | 3.1G | 11.7 | 188.7 |
|  | Canu | 16 | 24.58 | 30.67 | 4.09G | 37.58 | 118.57 |
|  | MECAT-CA | 16 | 4.36 | 4.23 | 10.05G | 5.87 | 14.46 |
|  | MECAT | 16 | 4.36 | 4.23 | 10.05G | 2.01 | 10.73 |
| D.melanogaster | FALCON | 16 | 64.20 | 45.21 | 10.26G | 31.31 | 140.72 |
|  | PBcR-MHAP | 16 | 62.09 | 28.57 | 3.40G | 9.5 | 101.22 |
|  | Canu | 16 | 12.70 | 28.37 | 4.01G | 27.01 | 69.34 |
|  | MECAT-CA | 16 | 2.98 | 5.75 | 10.29G | 10.47 | 19.20 |
|  | MECAT | 16 | 2.98 | 5.75 | 10.29G | 0.80 | 9.58 |
| Human | PBcR-MHAP(f) | 32 | 82000/32*1.6 |  | — | 33000 | 5750 |
|  | PBcR-MHAP(s) | 32 | 220000/32*1.6 |  | — | 40000 | 13000 |
|  | MECAT-CA | 32 | 231.11 | 24.12 | 80.77G | 817.75 | 1087.96 |
|  | MECAT | 32 | 231.11 | 24.12 | 80.77G | 60.93 | 230.54 |

All pipelines have been tested on the same computer with 2.0 GHz CPU and 512 GB RAM memory. For Human datasets, we cannot run PBcR-MHAP on our computer with 32 threads. Our computer is 1.6 times slower than the one used by Berlin et al. based on the running time of PBcR-MHAP using other data sets<sup>3</sup>. The running time of PBcR-MHAP is indirectly converted from those of Berlin et al<sup>3</sup>.

We further examine the quality of assemblies of five pipelines using four measures: assembly size, NG50^3,36^, number of contigs and the average number of contigs >200 bps per chromosome (ctg/chr) ^3,36^ (Table 8). In all five compared genomes, MECAT-CA and MECAT obtain comparable or improved assemblies. For *E. coli* K12, both MECAT-CA and MECAT recover the complete genome with just 1 contig. For *S. cerevisiae* W303^54^, both MECAT-CA and MECAT report close to perfect continuity with only 22 and 21 contigs, respectively. However, the MECAT obtain best NG50s with 100% assembly performance (even better than that of reference assembly S228C^55^), while MECAT-CA only report 89% assembly performance, which is similar to the results of PBcR-MHAP and Canu and better than the result of FALCON. For *Arabidopsis thaliana* Ler-0, MECAT reports only 56 contigs with better than reference assembly NG50 (100% assembly performance), which is much higher than those NG50 reported by other four pipelines with assembly performance from 68% of FALCON to 85% of PBcR-MHAP. For *D. melanogaster*, the MECAT also reports the highest NG50 with 82% assembly performance while it reports less total assembly size. For human CHM1, the assembly size reported by MECAT-CA is the closest to the size of human Ref 38 genome^56^, but it reported the largest number of contigs and largest ctg/chr ratio. Meanwhile, MECAT reported slightly better assembly size than those of PBcR-MHAP, but less than that of MECAT-CA. The NG50 reported by MECAT is 15% longer than those of PBcR-MHAP sensitive and MECAT-CA. And the number of contigs and ctg/chr reported by MECAT is similar to those of PBcR-MHAP sensitive, much less than those of MECAT-CA and PBcR-MHAP fast. Given its ability that MECAT can assemble the human CHM1^17^ genome in less than 10 days on a single 32-thread computer with comparable assembly quality, it can be an ultra-fast tool to assemble large genomes using SMS reads.

**Table 8.**
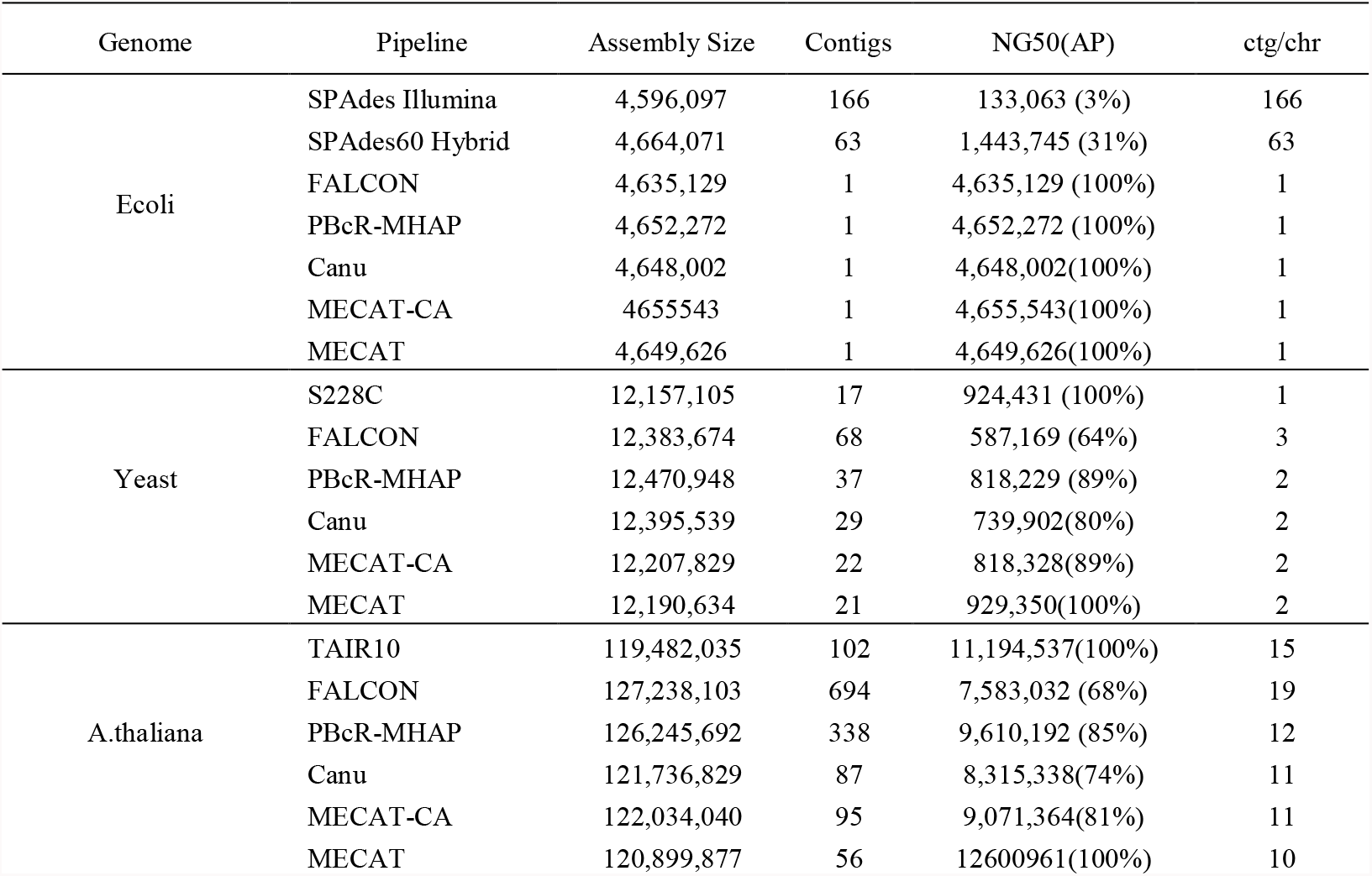
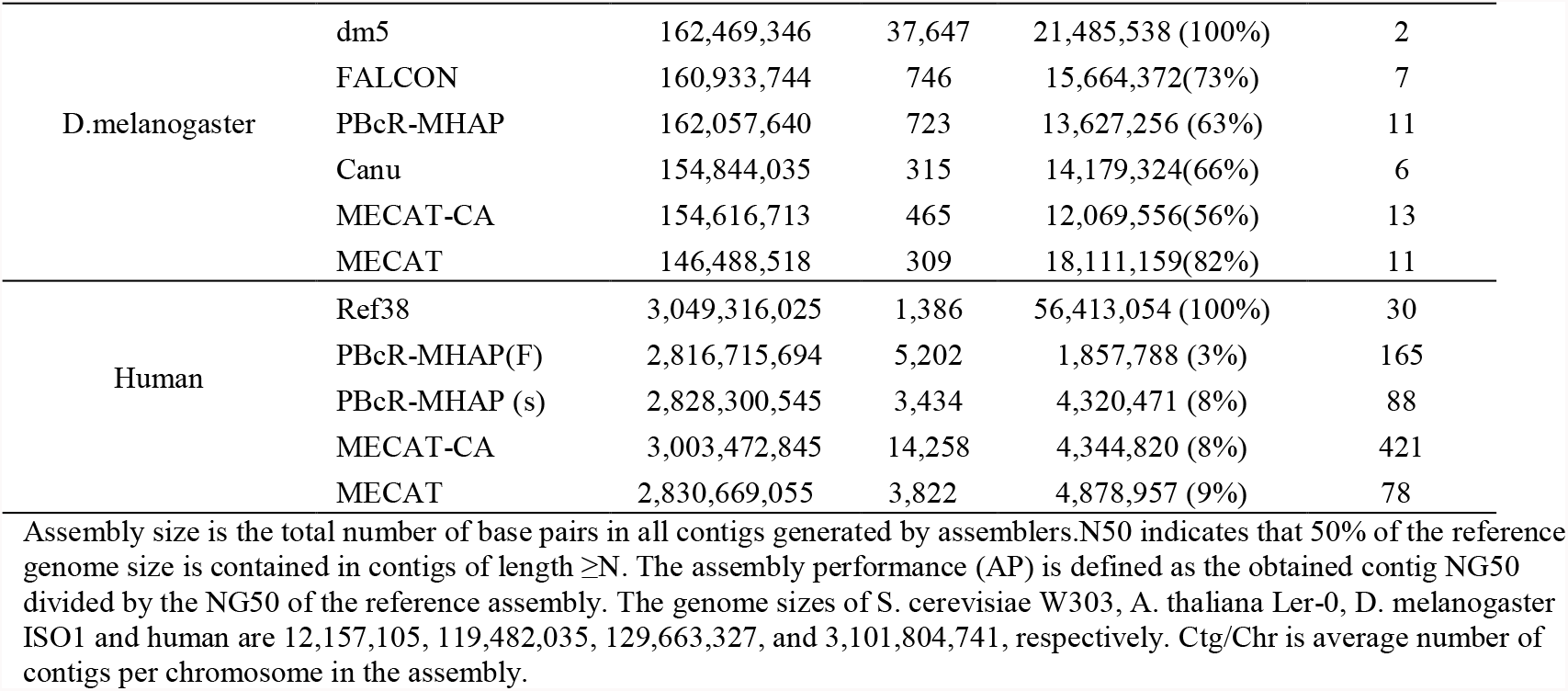
The assembly quality analysis of MECAT-Canu and MECAT.

| Genome | Pipeline | Assembly Size | Contigs | NG50(AP) | ctg/chr |
| --- | --- | --- | --- | --- | --- |
| Ecoli | SPAdes Illumina | 4,596,097 | 166 | 133,063 (3%) | 166 |
|  | SPAdes60 Hybrid | 4,664,071 | 63 | 1,443,745 (31%) | 63 |
|  | FALCON | 4,635,129 | 1 | 4,635,129 (100%) | 1 |
|  | PBcR-MHAP | 4,652,272 | 1 | 4,652,272 (100%) | 1 |
|  | Canu | 4,648,002 | 1 | 4,648,002(100%) | 1 |
|  | MECAT-CA | 4655543 | 1 | 4,655,543(100%) | 1 |
|  | MECAT | 4,649,626 | 1 | 4,649,626(100%) | 1 |
| Yeast | S228C | 12,157,105 | 17 | 924,431 (100%) | 1 |
|  | FALCON | 12,383,674 | 68 | 587,169 (64%) | 3 |
|  | PBcR-MHAP | 12,470,948 | 37 | 818,229 (89%) | 2 |
|  | Canu | 12,395,539 | 29 | 739,902(80%) | 2 |
|  | MECAT-CA | 12,207,829 | 22 | 818,328(89%) | 2 |
|  | MECAT | 12,190,634 | 21 | 929,350(100%) | 2 |
| A.thaliana | TAIR10 | 119,482,035 | 102 | 11,194,537(100%) | 15 |
|  | FALCON | 127,238,103 | 694 | 7,583,032 (68%) | 19 |
|  | PBcR-MHAP | 126,245,692 | 338 | 9,610,192 (85%) | 12 |
|  | Canu | 121,736,829 | 87 | 8,315,338(74%) | 11 |
|  | MECAT-CA | 122,034,040 | 95 | 9,071,364(81%) | 11 |
|  | MECAT | 120,899,877 | 56 | 12600961(100%) | 10 |
| D.melanogaster | dm5 | 162,469,346 | 37,647 | 21,485,538 (100%) | 2 |
|  | FALCON | 160,933,744 | 746 | 15,664,372(73%) | 7 |
|  | PBcR-MHAP | 162,057,640 | 723 | 13,627,256 (63%) | 11 |
|  | Canu | 154,844,035 | 315 | 14,179,324(66%) | 6 |
|  | MECAT-CA | 154,616,713 | 465 | 12,069,556(56%) | 13 |
|  | MECAT | 146,488,518 | 309 | 18,111,159(82%) | 11 |
| Human | Ref38 | 3,049,316,025 | 1,386 | 56,413,054 (100%) | 30 |
|  | PBcR-MHAP(F) | 2,816,715,694 | 5,202 | 1,857,788 (3%) | 165 |
|  | PBcR-MHAP (s) | 2,828,300,545 | 3,434 | 4,320,471 (8%) | 88 |
|  | MECAT-CA | 3,003,472,845 | 14,258 | 4,344,820 (8%) | 421 |
|  | MECAT | 2,830,669,055 | 3,822 | 4,878,957 (9%) | 78 |

Assembly size is the total number of base pairs in all contigs generated by assemblers. N50 indicates that 50% of the reference genome size is contained in contigs of length $\geq N$ . The assembly performance (AP) is defined as the obtained contig NG50 divided by the NG50 of the reference assembly. The genome sizes of *S. cerevisiae* W303, *A. thaliana* Ler-0, *D. melanogaster* ISO1 and human are 12,157,105, 119,482,035, 129,663,327, and 3,101,804,741, respectively. Ctg/Chr is average number of contigs per chromosome in the assembly.

### Validation analysis of assembly

We further validate the assemblies by comparing them to the reference genomes. Since MECAT is always faster and obtain better or comparable assembly performance, we only compared the assembled results of MECAT to those of PBcR-MHAH, Canu and FALCON. We first map the assemblies of *E coli*, Yeast, *Arabidopsis thaliana* and *D. melanogaster* to reference genomes using Nucmer^57^ (Supplementary Note 10), and then evaluate the mapping results using GAGE scripts^58^. Among four assemblies, only assemblies of *E coli* and *D. melanogaster* are generated from SMS read sampled from the same strains of reference genomes. All four assemblies are structurally consistent with reference genomes except some minor structural variation (Supplementary Figs. 6-22). Supplementary Table 3 provides GAGE^58^ accuracy metrics for these assemblies. With all discrepancies between assembly and reference genome sequence being counted as error, the assemblies reported by MECAT are still at least 99.99% accuracy (QV=40^3^) compared to the reference genomes. We also align four assemblies before and after Quiver^53^ polishing onto reference genome using dnadiff program^52^, and count the single-nucleotide polymorphisms (SNPs) and big indels (>10bps). The numbers of SNP and indels in assemblies reported by all four pipelines are similar, especially after Quiver polishing (Supplemental Table 3). We further map all 17294 annotated genes of D. melanogaster^59,60^ onto the assemblies. We identify a total of 16972, 17044, 17055 and 16839 genes mapped to a single contig in a single alignment from assemblies of PBcR-MHAP, FALCON, Canu and MECAT, respectively, while 16944, 17020, 17037 and 16812 genes of these have over 99% identity. The results show that the qualities of assemblies from MECAT are comparable to those from other pipelines.

Solving the repeat regions is the most import task in genome assembly. We evaluate four assemblies of *D. melanogaster* by comparing the completeness of transposable element (TE) families^61^ (Supplementary Note 12 and Supplemental Table 4). In all 5,433 annotated TEs from the flybase^3^. MECAT assembly contains 5301 (97.6%) TEs, in which 5141 (94.6%) aligned perfectly to the reference. Meanwhile, PBcR-MHAP, FALCON, Canu assemblies contain 5274, 5306 and 5319, respectively. And 4984, 5190 and 5165 of them are aligned perfectly. We have examined two TE families, *roo* and *juan*, in detail. In MECAT assembly, 131 of 138 copies in *roo* TE family are aligned. Of these, 123 are perfectly aligned. For 11 elements of the *juan* family, all are perfectly confirmed. Those results are similar to the assemblies of other three pipelines (Supplementary Table 4). The TE analysis in *D. melanogaster* demonstrates that MECAT is capable of reconstructing TE repeats sequences accurately.

To further evaluate the ability of MECAT that reconstructs the repeat regions of genomes, we have examined the telomeric repeats in *S. cerevisiae* assembly of MECAT (Supplementary Note 13 and Supplementary Table 5). We are able to map telomeric repeats of 15 chromosomes to assembled contigs. Among them, seven chromosomes assembled in a single contig have at least 50% of terminal telomeric repeats mapped on both ends. For chromosomes contains more than one contig, we have mapped the two end telomeric repeats onto two contigs of six chromosomes and one end telomeric repeats onto one contig of one chromosome. There are telomeric repeats of two chromosomes cannot be mapped onto any contig, which may due to either assembly error or different strains. Our results are similar to those obtained from assemblies of PBcR-MHAP, FALCON and Canu (Supplemental Table 3-5).

### De novo assembly of a human diploid genome

Finally, to demonstrate MECAT in *de novo* assembly of large genome, we have assembled 102x SMRT sequencing reads from a diploid Han Chinese genome using MECAT on a 32-core computer. It takes only 25 days to finish the whole assembly. The Han assembly is submitted and assessed by NCBI (https://www.ncbi.nlm.nih.gov/assembly/GCA_001856745.1/). We compare our Han assembly to another Han Chinese genome assembly from BGI, YH1, which is assembled from Illumina data (http://yh.genomics.org.cn/). The total size of our Han assembly is 2,908,568,123bp, which is much longer than the 2.2G bp of YH1. The NG50 of Han assembly is 8,583,694 and is 2263 times longer than NG50 of YH1. The Han assembly only has 4456 contigs and is only 0.6% of total number of contigs of YH1. Furthermore, the Han assembly has six contigs with size greater than 40M. The length of longest contig of Han is 66M and is much greater than the 0.9M of YH1. Moreover, the Han assembly shows much better continuity of genome comparing to the YH1 assembly (Figure 3). Thus, our assembly can be a better reference genome for Han Chinese and it also show that SMRT sequencing can significantly improve de novo assembly quality and integrity.

**Figure 3.**
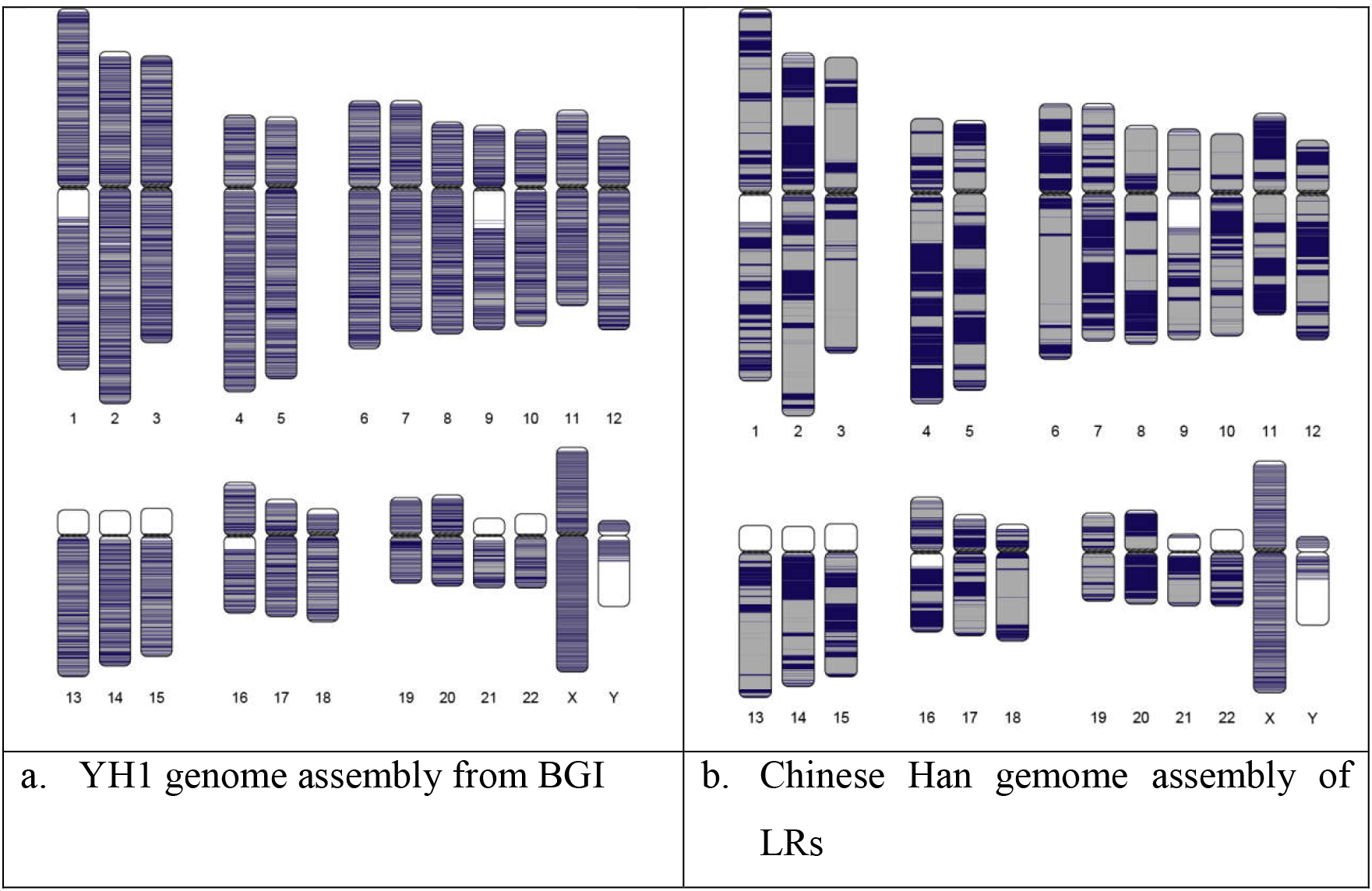
Comparison of the continuity of two Chinese assemblies. We paint the assembled contigs on human chromosomes using the colored Chromosomes package. The black and gray shades indicate contigs and transitions between shades indicate contig boundary or alignment breakpoint. White regions indicate missing assembly sequence or uncharacterized reference sequence with no contig mapped. A) the Illumina-based YH1 assembly from BGI. B) the Han assembly by MECAT.

We also map the Han assembly onto hg19 human reference genome using Nummer software and the dotplot figure (Supplementary Note 14 and Supplementary Fig. 25) shows that our assembly is structurally consistent with hg19 genome except for some minor structural variation. Furthermore, we have aligned the Illumina datasets from YH1 onto Han, YH1 assemblies and hg19 human reference genome using bowtie2^50^ (Supplementary Note 14 and Supplementary Fig. 24). The Han assembly gains the best mapping rate (83.81%) comparing to the mapping rates to YH1 (73.06%) and hg19 (82.05%). This result validates that Han assembly is a better reference genome.

To find structural variations between Han Chinese genome and European genome, we have mapped both Han and YH1 assemblies onto hg19 human reference genome. We find 29836 structural different genome region (≥ 20bp) in Han assembly and most of them (65%) can also be identified in YH1 assembly (Supplementary Table 6). We have also compared Han assembly to the Korean genome^62^. The result shows that the Han assembly is structurally consistent with Korean genome except minor structural variants (Supplementary Figs. 28). We have further compared MHC region of Han assembly with those of Korean and hg19 genomes. The result (Supplementary table 6) shows that Han and Korean assemblies have 157 and 147 variants (>10bp) comparing to hg19 and 50 of them are the same site. Among two recently validated max. variants (54896bp and 10286bp) in MHC region of Korean 62, only one variant (10286bp) has been found in Han assembly. This results show that there are structural different between the MHC regions in Han Chinese and Korean although 33% of those regions are the same.

## Discussion

The repeat regions in the genome lead to the higher frequency of k-mer mapping between SMS reads. When k-mer matching is used to filter random pairs, all-pair alignment may produce excessive candidate pairs, and local alignment of excessive candidate pairs consume the major computational time in SMS read correction. Thus, reducing the redundant repetitive k-mer matches is the key to reduce excessive candidate pairs, and then computational cost. However, completely masking low-complexisty sequence or ignoring highly repetitive k-mer may lead to the lost of some correct overlaps^42^. Recently, the Canu pipeline employed a tf-idf k-mer weighting method to reduce the effects of repetitive k-mer matches. However, even with k-mer weighting, the MinHap algorithm in Canu only report the local matched k-mers pairs without considering the arrangement of k-mer pairs, which may still lead to excessive matches. In BLASR, the best arrangement of k-mer pairs is solved by slow sparse dynamic programing. Here, our scoring algorithm considers matched k-mer pairs between two reads as well as intervals between matched k-mer pairs. Furthermore, the repetitive matched k-mer pairs are removed from scoring, which reduces the effect of small repetitive region in read overlapping. Our algorithm provides a heuristic global alignment score between two reads, which is more sensitive to the true overlap. One proven of this is that the MECAT can be used to align SMS reads to reference genome with high sensitivity and precision similar to BLASR

Another benefit of our global alignment score is selecting the top informative matched reads for a give read template. Since the top informative matched reads selected by the global score are so reliable, we even do not need to perform local alignment using diff program^46^ to further filter them, which has reduced the computational cost for the whole read correction step.

The alignment tool in MECAT can also be plugged into other pipelines. For example, the Celera Assembler (CA^35^), an overlap-layout-consensus (OLC) based assembler, need the overlap length between the reads to obtain high quality assemblies. Since our alignment score is correlated with the overlap size between two reads. Thus, we can replace the overlapInCore in Canu^42^ (v1.0) (a new version of CA for SMS reads), which uses a slow blast-like algorithm for computing overlaps of corrected reads, with alignment tool in MECAT. This allows us to dramatically reduce the computational time for contig construction.

With the new alignment algorithm as well as the improved read correction method, we are able to develop a new assembler pipeline, MECAT, which is capable to produce high quality *de novo* assembly of large genome from long noisy SMRT reads with low computational cost. Our experimental results show that MECAT can assemble a high quality human genome assembly using 54X SMRT reads in only 7600 CPU core hours and a high quality Chinese Han human assembly using 102X SMRT reads in 19200 CPU core hours, which is ten times faster than current fastest assembler. The MECAT makes it possible to assembly large genomes on a single server computer and small clusters using SMRT reads.

Currently, the structurally divergent alleles are not considered in the MECAT pipeline. One of our future works to improve MECAT is designing new algorithm to distinguish structurally divergent alleles, and then make MECAT able to assemble polyploidy genomes. In this paper, we focus on assembling genomes using PacBio SMRT reads. Since Oxford Nanopore reads have similar characteristics as SMRT reads, we will evaluate the applicability of MECAT to Nanopore reads in the future.

## Methods

### Indexing and matching of reads

The finding of potential matching between reads is based on the matching of k-mers (substrings with length of k) in the reads. A read *r* of length L has total L-k+1 k-mers. We first index the reads using a hash table with the k-mers as key. We consider the overlapping k-mers between the blocks of reads. For each read, we break it into multiple blocks with each block of length *B*, which is usually be 1,000 bp or 2,000 bp. The values in the hash table are the position of k-mers in the blocks of reads.

To search for the matching reads, we scan the k-mers in blocks of reads and look up the matchings in the hash table. We break the reads into blocks of same length *B*. In order to reduce the computer time, we only sample the k-mers in each searching block. We slide a k-sized window along each block with a step length of *sl*. Thus, the number of k-mers in the search is only about 1/*sl* of the number of total k-mers from reads. A typical value of *sl* is 10. A searching block is matched to an indexing block if the number of their overlapping k-mers is greater than a predefined threshold *m*. Two reads are considered as matched if at least a pair of blocks is matched between them.

Given two read blocks of length *B*, the number of k-mers sampled from the search block is (B/sl-1). Let *O* be the overlapping length of a pair of matched blocks, *O*≤*B*. The expected number of matched k-mers in O is^3^

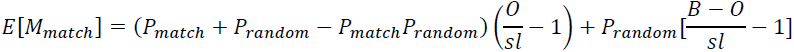

where, *P_random_* is the probability of a random k-mer and *P_match_* is the probability that two k-mers are matched. As the block length *B* is fixed, for a given error rate and no repetitive sequence, the number of matched k-mers between two blocks grows with the overlapping length *O*. For a highly matched block pair, *P_match_* >>*P_random_*. The expected number can roughly be estimated as:

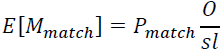

### Filtering false matched reads using global score

We develop a new pseudo linear global scoring algorithm to filter the excessive, non-informative matched reads. Our scoring algorithm has two steps. The first is the mutual scoring step. For each matched read pair, we first randomly select a matched block pair and mark it. Then, we score the matched k-mer pairs in this matched block pair. Let (p_i_, p_j_) be the positions of *i*-th and *j*-th k-mer in one block and (p′_i_, p′_j_) be the position of *i*-th and *j*-th k-mer in another block of matched pair. We define the distance difference factor (*DDF_i,j_*) between *i*-th and *j*-th k-mer as following:

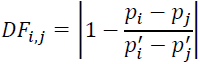

If *DF_i,j_<ε*, which indicates that the both k-mers supporting each other, we increase the scores of both k-mers by 1. The *ε* is set to 0.3 in our experiments. By calculating the DF between all possible pair of k-mers, we obtained scores for all overlapping k-mers of matched blocks. We only use the non-repetitive k-mer pairs in our scoring. If a k-mer is matched more than once in the same block, it will be excluded from scoring. If the score of k-mer with highest score is significant (greater than a threshold), we set it as the seed position for future alignment. If there are multiple k-mers having the same score, we randomly select one as seed.

The second is the extension scoring step. In order to increase the reliability of seed and reduce the computation of whole scoring process, we extend the scoring process from selected block pair to its neighbor matched block pairs if a seed k-mer is obtained. For each overlapping k-mer in neighbor block pair, we calculate the DF between the k-mer and the seed k-mer in original block pair. If *DF*<*ε*, we increase the score of seed k-mer by 1. If 80% of DF values of overlapping k-mers in a neighbor block pair satisfy *DF*<*ε*, we mark the block and do not score the k-mers in this block pair in the future. After one loop of mutual and extension scoring process, if there are still matched block pairs having not been marked, we continue the scoring process on those block pairs. The mutual scoring is done in O(N^2^) time and the extension scoring is done in O(N) time, where N is the number of k-mer matches As the number of k-mers in mutual scoring is small, the overall scoring process can be done in a pseudo linear time.

### Aligning SMS reads

The pair alignment of SMS reads is the first step for correcting reads and assembling genome using SMS reads. We only consider two SMS reads have more than 2000bp overlapped. Namely, we set the block length to 2000bp. After scoring the matched k-mers between two SMS reads, we sort the k-mers based on their scores. Then, we use the top ranked k-mers as seeds to perform local alignment of two reads using diff algorithm^40,46^. If the overlapped length between two SMS reads is longer than 2000 bp and the mismatch rate of overlapped sequence is less than twice of SMS read error rate, we consider there is a match and output the align results. All detail parameters are described in Supplementary Note 2.

### Aligning SMS reads to a reference genome

The procedure of aligning SMS reads to a reference genome is similar to those of aligning two reads. We index the reference genome sequence and search the reads from the index table. We first break the reference genome into blocks with length of *B* and index the k-mers in each block. Then, we break the reads into blocks of same length *B* and sample the k-mers for looking up the index table. The matched k-mers between a read and reference genome are also scored. The top ranked k-mers are used as seeds to perform further local alignment using diff.

In order to obtain high sensitivity of the alignment of SMS reads to a reference genome and keep the computational cost low, we present a two-steps approach. In first step, we use the block length *B* of 1000bp and k-mer sampling step length *sl* of 20 to align reads to reference genome. As some SMS reads have less match k-mer or the distribution of their matched k-mers is uneven, those SMS reads cannot find the matching position in first step. In the second step, we double the block length *B* to 2000bp and half the k-mer sample step length *sl* to 10, and align so far unmatched reads again. In practice, we can find the matching for significant amount of reads in first step and can find the matching for most of reads after second step. As the computational cost of second step is much higher than that of first step, our two-steps approach allows us reduce the computational cost and maintain high sensitivity at the same time. All detail parameters are described in Supplementary Note 1.

### Correcting SMS reads

The random and independent properties of errors in SMS reads make it possible to correct them. Generally, there are two steps to correct SMS reads. The first step is building a multiple read alignment for each read to be corrected. The second step is constructing the correct read from the consensus of alignment. For the first step, we use our own alignment tool. We perform pairwise alignment (without local alignment using diff) between all reads with length greater than 5000 bp. Then, we filter out alignment if the overlapped sequence is less than 90% of shorter read in the pair. The output of the alignment is written into multiple files. Each file includes the alignment information of 200,000 reads. For the second step, we develop a new SMS read error correction method by combining the principles from both DAGCon and FalconSense. We summarize the pairwise alignment to construct a consensus table with the counts of match, insertion and deletion. For trivial regions with consistent matches: match_count/(match_count+deletion_count)>0.8 and no significant insertion occurring (insertion_count<6), or consistent deletions: deletion_count/(match_count+deletion_count)>0.8 and no significant insertion occurring (insertion_count<6), we can simply determine the consensus base according to the count. For other complicate regions, we construct a local POG and solve the consensus with dynamic programming. All detail of this algorithm is described in Supplementary Note 2.

### *De novo* assembly using SMS reads

We develop a new pipeline for assembling SMS reads by integrating our new alignment and error correction method with the Celera Assembler (CA). Our pipeline has three steps. In first step, for each reads longer than 3000bp, we perform pairwise alignment against other reads and select 100 matched reads with top matched scores. No detail local alignment is performed in this step. In second step, we correct all template reads (>3000bp) using their matched reads. Finally, we pairwise align the corrected reads using our alignment method and feed the results into the “Unitig Construction” module of CA or Canu to construct the unitigs. We can also feed our correct reads to the CA directly, which used the overlapInCore for pairwise alignment.

### Evaluation

We evaluate MECAT tools using both simulated reads and raw reads from five model organisms. We compare our alignment tool to previous pairwise alignment tools, including BLASR, MHAP and Daligner, and to genome alignment tools, including BLASR and BWA-mem. We compare our correction tools to those in PBcR and Falcon. We also systematically evaluate our assembly tools by comparing with Canu, PBcR-MHAP, Falcon. The details of those comparisons are reported in Supplemental Note 5-9.

### Accession codes

Assembly and annotation files of Han-1 Chinese Human are available from GenBank: GCA_001856745.1. All source codes for MECAT and the analyses presented here are available from https://github.com/xiaochuanle/MECAT. The software and data used for this manuscript (including supplementary files and scripts) are available from http://sysbio.sysu.edu.cn/software/MECAT. Note: Any Supplementary Information and Source Data files are available in the online version of our paper.

## ACKNOWLEDGMENTS

We are grateful to De-Pei Wang for supplying Chinese human Dataset and some advices throughout this project. We thank NCBI assembly group for Han-1 Chinese annotation. Here, We also thank Pacific Biosciences and all those involved in generating and freely releasing the data analyzed here. This work was collectively supported by National Natural Science Foundation of China (31471789, 31600667 and 31200612), the Fundamental Research Funds for the Central Universities (15ykjc23d) and Guangdong Natural Science Foundation (2015A030313127).

## AUTHOR CONTRIBUTIONS

C.L.X. conceived and designed this project. C.L.X. conceived, designed and implemented alignment algorithm. Y.C. and C.L.X. conceived, designed and implemented the consensus algorithm. Y.C. and C.L.X integrate all programs into MECAT pipeline and the documentation writing. S,Q.X. C.L.X. and Y.C. ran and analyzed the genome assemblies and our algorithms performance. K.N.C. and Y.W. coordinated data release and assisted with pipeline executions. C.L.X and Y.C. drafted the manuscript. C.L.X. and S.Q.X. drafted supplementary files and the analysis scripts of results. F.L. provided the theoretical analysis of algorithms in this paper. F.L. rewrote and improved all manuscripts and Z.X. revised supplementary files and some sections in paper. All authors read and approved the final manuscript.

